# Long-term daily light exposure boosts photoreceptor maturation in retinal organoids

**DOI:** 10.1101/2025.07.17.665292

**Authors:** E. M. van Oosten, A. D. M. Hoogendoorn, W. Kieboom, S. L. Berendsen, J. Haerkens, M. M. E. van der Maden, D. Veerman, A. Bufe, S. Almedawar, A. D. van der Meer, T. Barta, R. W. J. Collin, A. Garanto

## Abstract

Induced pluripotent stem cell-derived retinal organoids (ROs) have become promising personalized models to study inherited retinal diseases (IRDs) and develop new innovative therapies. Although ROs mimic key retinal features and show some light responsiveness, their differentiation and maturation remain limited and lengthy. We hypothesize that standard dark culture conditions limit the expression of genes and proteins related to retinal function, delaying differentiation and complicating disease modeling. Therefore, we investigated whether daily light exposure could promote photoreceptor maturation. ROs were exposed to six hours of daily light starting from day 70 *in vitro*. Light conditioning led to enhanced photoreceptor maturation, specifically an increase in rod photoreceptors, a late-born cell type, without signs of increased stress or cell death. Our findings suggest that daily light exposure enhances RO differentiation, opening new avenues to investigate molecular and cellular phenotypes in IRDs and accelerate therapy development in more relevant functional models.

## Introduction

Inherited and multifactorial retinal diseases pose a major burden on patients and society, yet effective treatment options are limited. One major limitation in therapy development is the poor translation of preclinical results into clinical trials, as there are substantial differences between the preclinical models and the human eye.^1–3^ Commonly used *in vivo* models do not always recapitulate human disease ^4,5^, and most *in vitro* models are too simplistic and lack the three-dimensional retinal microenvironment.^6,7^

In recent years, induced pluripotent stem cell (iPSC)-derived cell models have gained popularity for disease modeling and therapy development.^8,9^ iPSC-derived retinal organoids (ROs) self-organize into a distinct layered structure resembling the human retina by mimicking human retinal development in a dish.^10^ ROs consist of an outer nuclear layer containing rod and cone photoreceptors, and inner layers containing other retinal cell types including retinal ganglion cells and bipolar cells. These ROs are currently being used to model retinal diseases and have been proven effective in testing therapies in a human context^9,11–13^

Despite these advances, ROs remain developmentally immature. Specifically, single-cell RNA-sequencing experiments show that ROs more closely resemble fetal retinal tissue.^14,15^ This developmental immaturity limits retinal disease modelling, as most inherited retinal diseases (IRDs) occur in childhood or early adulthood, instead of in fetal tissue.^16^

Although ongoing efforts to improve RO culture times seem promising, for example by perfusion or vascularization of ROs, alternatives are needed to further enhance maturation.^17,18^ Typically, ROs are cultured in a dark incubator and are only taken out of the incubator for media refreshment and monitoring; about three times per week for long-term cultures. The lack of light that ROs are exposed to might influence photoreceptor maturation, as it is known that dark-reared mice have disrupted photoreceptor responses.^19–21^ Additionally, many of the photoreceptor transduction genes and proteins undergo changes in expression, morphology, and localization when exposed to light.^22,23^ Without light exposure, these changes in gene and protein expression, morphology, and localization are unlikely to occur, which could limit the maturation of photoreceptor cells and the corresponding functional readouts for IRDs.

In this study, we tested whether long-term, daily light exposure could promote photoreceptor maturation in ROs. For that, we cultured ROs in a daily light cycle regime from 70 days *in vitro* (DIV70) onwards. We then assessed the expression of photoreceptor cell markers to determine whether light-conditioned ROs exhibited enhanced photoreceptor maturation compared to control ROs.

## Results

### Daily 6-hour light stimulation enhances photoreceptor gene expression in retinal organoids after 5 weeks

We hypothesized that the lack of light during RO culture hinders the differentiation and maturation of photoreceptors. To address this, we generated ROs and exposed them to daily light cycles (350-400 LUX, constant white light) (**Figure 1a-c**). Initial experiments showed that when ROs were exposed to light before DIV70, the organoids were dying and disintegrating, while when starting at DIV70 organoids survived and developed properly. Therefore, we applied cycles of 6 hours of daily light and 18 hours of darkness from DIV70 onwards and assessed gene expression after five weeks in this routine (**Figure 1d**). Six hours of daily light increased the expression of photoreceptor markers after five weeks of culturing (DIV105), compared to ROs that were cultured completely in the dark (**Figure 1e**). This includes a significant 1.66-fold increase in the expression of the transcription factor *CRX*. As a likely consequence, we found upregulation of photoreceptor inner segment (IS) and outer segment (OS) markers *ARR3*, *PRPH2*, and *RCVRN* indicating increased differentiation of photoreceptors in light-conditioned ROs. We also investigated if cycles of 12 hours light/dark would be more effective. Surprisingly, when exposing ROs to 12 hours of daily light, we observed a decreased expression of photoreceptor markers, indicating a detrimental effect on the ROs in our set-up (**Supplementary Figure 1a**). We therefore decided to continue the rest of the experiments using six hours of light on a daily basis starting at DIV70.

**Figure 1.**
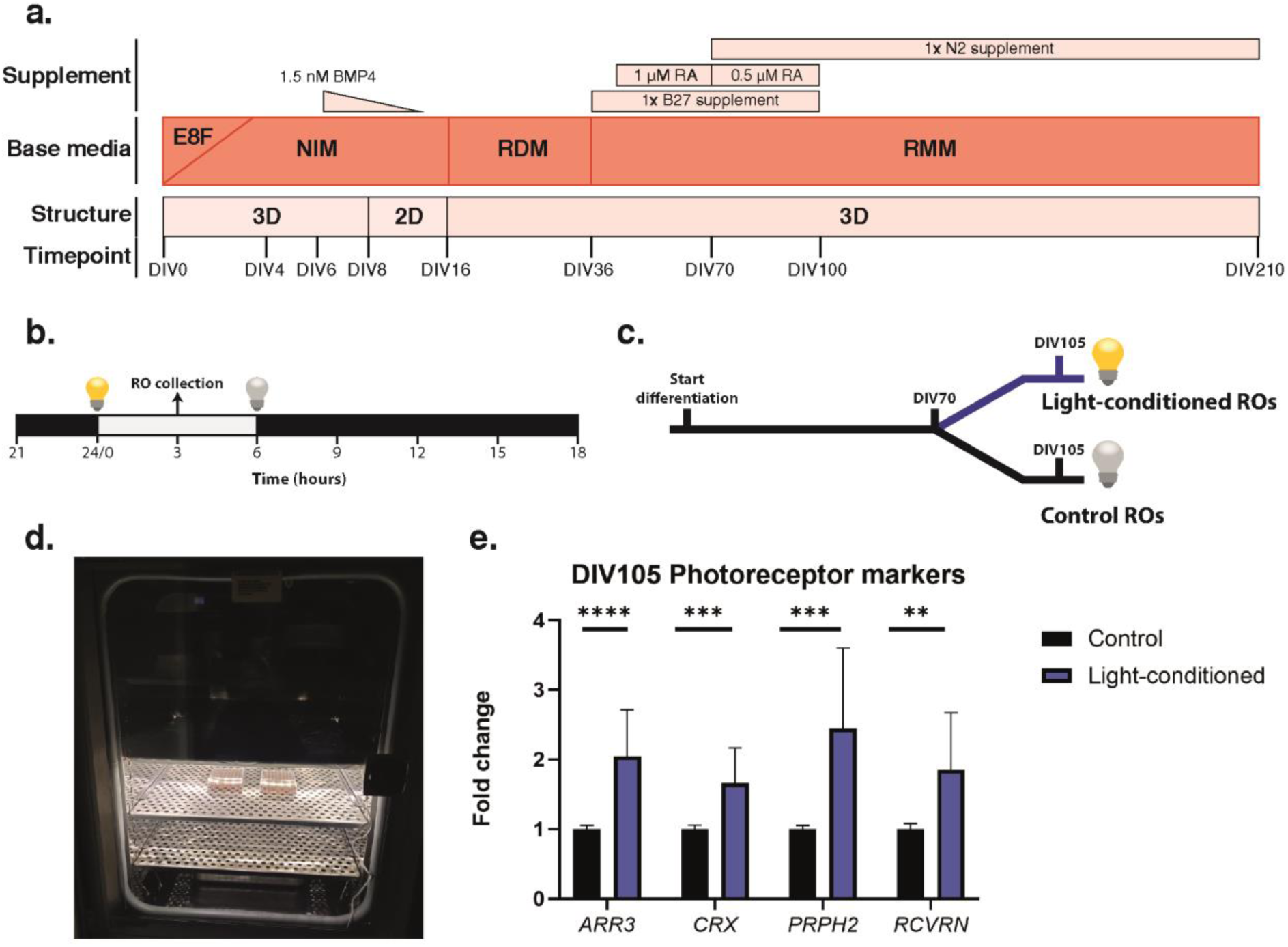
Experimental setup for studying the effect of light conditioning on RO differentiation. **(a)** Schematic representation of the RO differentiation protocol. **(b)** Schematic representation of the light-dark cycle set up at 350-400 LUX during 6 hours on a daily basis. ROs were collected in the middle of the light cycle for all experiments. **(c)** Diagram depicting the experimental design in which light was applied from DIV70 until DIV105. **(d)** Photograph of the technical setup in the incubator in which organoids were exposed to light or cultured in regular dark condition. **(e)** Gene expression analysis of photoreceptor markers from four independent RO differentiations with three technical replicates each. Values were normalized against *GUSB* expression and represented as the mean ± SD. Statistical differences are indicated with asterisk (** p-value<0.01; *** p-value<0.001; ****p-value<0.0001) using an unpaired t-test.

### Long-term daily light exposure results in increased photoreceptor markers

To assess whether the positive effects on photoreceptor gene expression persist over time, we extended the light conditioning to DIV210 of two independent retinal organoid differentiation batches, while collecting ROs for qPCR every five weeks (DIV105, DIV140, DIV175, DIV210) (**Figure 2a**). All ROs that remained in culture up to DIV210 were imaged one week prior to harvesting (DIV202) with phase-contrast microscopy to measure photoreceptor OS length. In total, 18 control and 40 light-conditioned ROs were analyzed. The OS length was measured in five photoreceptor OSs per RO using FIJI and revealed an increased OS length in light-conditioned ROs in our set-up (**Figure 2b**). To determine whether photoreceptor gene expression was improved in light-conditioned retinal organoids across timepoints, we calculated the fold change versus the baseline (DIV70) gene expression of each differentiation batch. In both control and light-conditioned ROs gene expression of major photoreceptor markers (*CRX, PRPH2, PDE6H, RCVRN, RHO,* and *ARR3*) increased over time compared to DIV70 (**Figure 2c**). We also detected trends in increased photoreceptor gene expression in light-conditioned ROs compared to control ROs during maturation stages DIV140 and DIV175, albeit with some variability. We found that specifically at the last timepoint, DIV210, light conditioning significantly showed an increase in photoreceptor gene markers *CRX, PRPH2, PDE6H, RCVRN, RHO,* and *ARR3,* suggesting a more advanced maturation.

**Figure 2.**
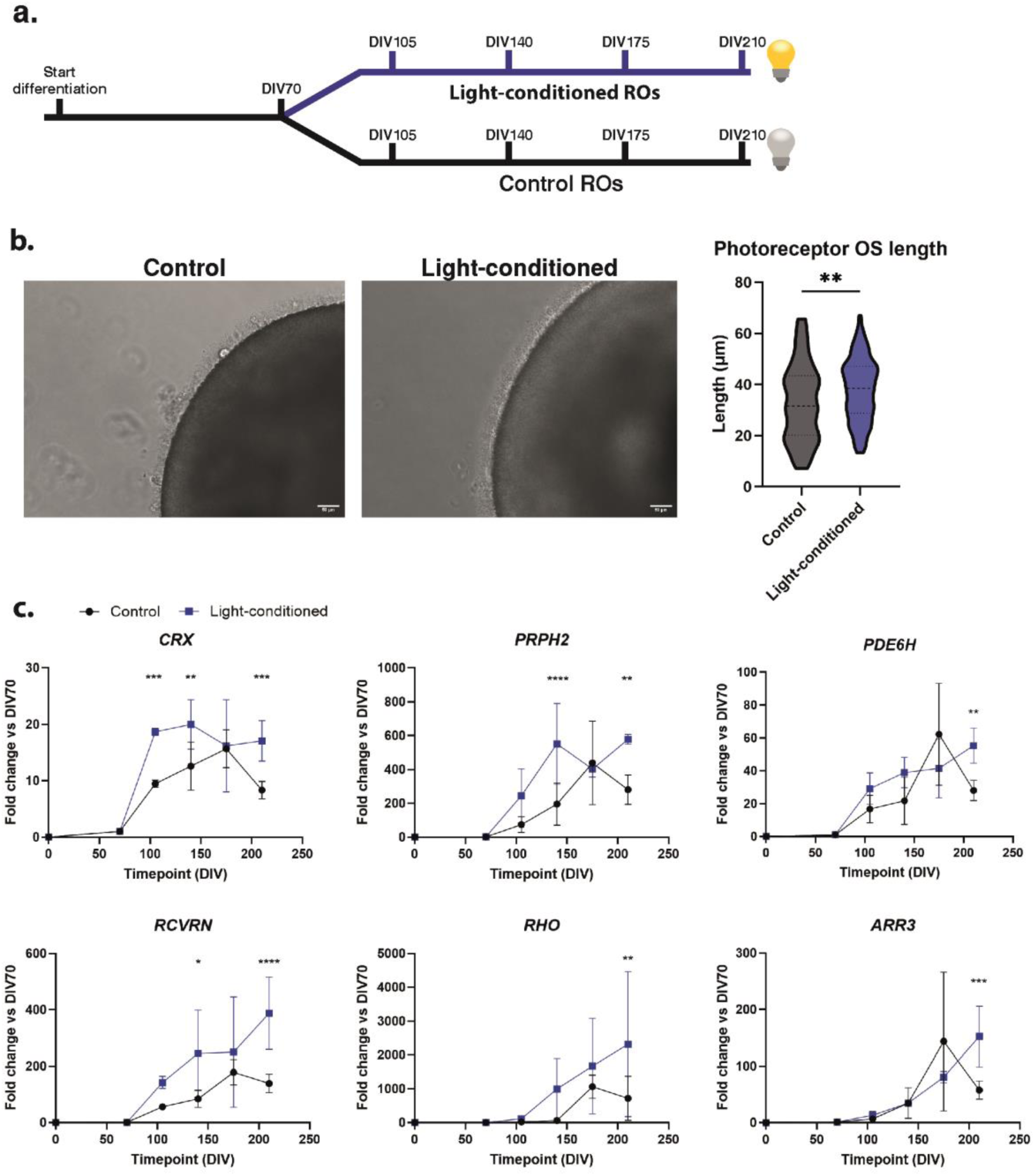
Daily light exposure enhances photoreceptor outer segment generation and marker expression in retinal organoids. **(a)** Schematic representation of the experimental protocol. ROs were split into control group and light-conditioned group from DIV70 on, and were collected for gene expression analysis every five weeks. **(b)** Representative images of control and light-conditioned ROs at DIV202. The violin plot represents the quantification of the outer segment length. Statistical differences are indicated with asterisks (** p-value<0.01) and were calculated using Mann-Whitney test. Scale bar = 50 µm. **(c)** Gene expression analysis of photoreceptor markers in control (black) and light-conditioned (blue) ROs at different time points compared to DIV70. Values are represented as the fold change mean ± SD from two independent differentiation batches with three technical replicates each. Statistical differences are indicated with asterisks (*p-value<0.05; ** p-value<0.01; *** p-value<0.001; ****p-value<0.0001) and were calculated with two-way ANOVA followed by a post-hoc test.

### Proteomic analysis of light-conditioned retinal organoids

To investigate whether the increased photoreceptor gene expression found at DIV210 correlates with increased protein levels, we performed a proteomic analysis of control and light-conditioned ROs. We analysed four samples containing three retinal organoids at DIV210 (control and light-conditioned) for proteomic analysis. After data collection, proteins that were detected in two or fewer samples per condition were excluded to ensure reliability of the results. Principal component analysis revealed that, for both conditions, three out of the four samples clustered together, indicating sample-to-sample variability (**Supplementary Figure 2**). Differential expression analysis of data revealed significantly enriched and depleted proteins in light-conditioned ROs (**Figure 3a**). Only 30 upregulated and 26 downregulated proteins had a p-value < 0.05 and a log2FC ≥ 0.5, as indicated in the volcano plot (**Figure 3b**). Gene Ontology (GO) analysis for biological processes of these proteins resulted in a few differentially regulated pathways between conditions related to metabolism (**Supplementary Figure 3a**). In addition, we analyzed all significantly up- or downregulated proteins regardless of the fold change to find disrupted pathways. Here, we mainly detected pathways related to ribosome assembly and biogenesis (**Supplementary Figure 3b**). Given the relatively small number of differentially expressed proteins, and therefore limited changes in pathways, we decided to proceed with data analysis by looking at photoreceptor-specific proteins to assess photoreceptor differentiation.

**Figure 3.**
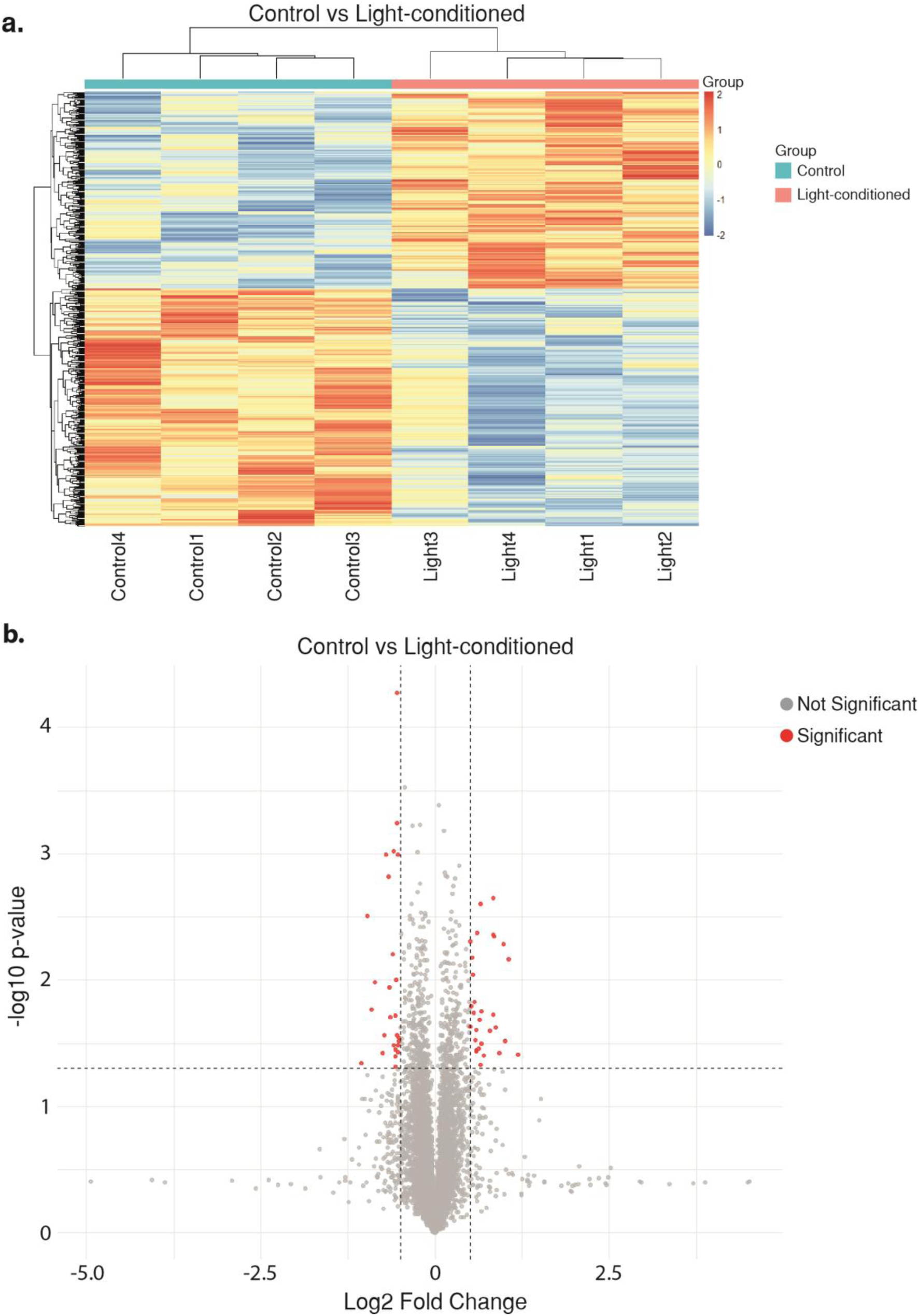
Light-conditioned ROs show differentially expressed proteins compared to controls at DIV210. **(a)** Hierarchical clustering heatmap displaying normalized expression values (z-scores) of differentially expressed proteins between control (blue) and light-conditioned (pink) samples. Each row represents a protein, and each column represents an individual sample (n = 4 per group). Expression levels are color-coded: red indicates higher expression, blue indicates lower expression. Clustering was performed on both protein (rows) and samples (columns), highlighting distinct patterns between the two experimental groups. **(B)** Volcano plot of differential protein expression between control and light-conditioned ROs. The plot describes the distribution of proteins based on their log2 fold change (x-axis) and statistical significance (-log10 p-value, y-axis). Each dot represents a protein. Significantly differentially expressed proteins are highlighted in red. Vertical dashed line indicates the fold change threshold, and the horizontal dashed line marks the significance threshold (p < 0.05).

### Light conditioning of ROs results in increased rod photoreceptor protein expression at DIV210

We aimed to determine whether light conditioning increased the expression of photoreceptor proteins. We therefore compiled a list of proteins based on gene ontology (GO) annotations, associated with photoreceptor inner segments (GO:0001917) and photoreceptor outer segments (GO:0001750). We then plotted the log2FC of each protein, and found both proteins that were enriched, as well as reduced in expression (**Supplementary Figure 4a**). Here, we noticed that rod-associated proteins were often enriched in expression. As a next step, we generated a list of detected proteins of either cone photoreceptor-related (ABCA4, ARR3, CNGB3, GNAT2, GNGT2, PDE6C, PDE6H) and rod photoreceptor-related (CNGB1, GNAT1, GNGT1, NR2E3, PDE6A, PDE6B, PROM1, PRPH2, RHO, ROM1, SAG) markers. Interestingly, light-conditioned ROs have increased expression of rod-specific photoreceptor markers, while cone-specific photoreceptor markers were either unchanged, or show a trend towards a reduction (**Figure 4a**). To validate these findings, we performed western blot analysis on the same samples. Since ROs are heterogenous both in size and content, we considered that TUBG1 was only providing information on the total amount of protein but not the number of specific cell types. To account for the number of photoreceptor cells, the cells of interest, we normalized the expression of ABCA4 and RHO to CRX, and calculated fold changes relative to control samples. Again, we were able to detect an increase in the expression of both ABCA4 and RHO in light-conditioned ROs (**Figure 4b**). The increases in especially rod-specific photoreceptor markers might indicate that the maturation of ROs is accelerated in light-conditioned ROs, favoring the differentiation of late-born rod photoreceptors. With immunofluorescent labelling of DIV210 control and light-conditioned ROs for pan-photoreceptor marker CRX and rod-specific transcription factor NRL, we were able to label both cone nuclei (CRX+, NRL-) and rod nuclei (CRX+, NRL+) in our ROs (**Figure 4c**). Overall, we show that light-conditioned ROs have increased photoreceptor protein expression and increased appearance of rod versus cone photoreceptors, indicating that light conditioning has an impact on the photoreceptor fate, potentially leading to more mature ROs.

**Figure 4.**
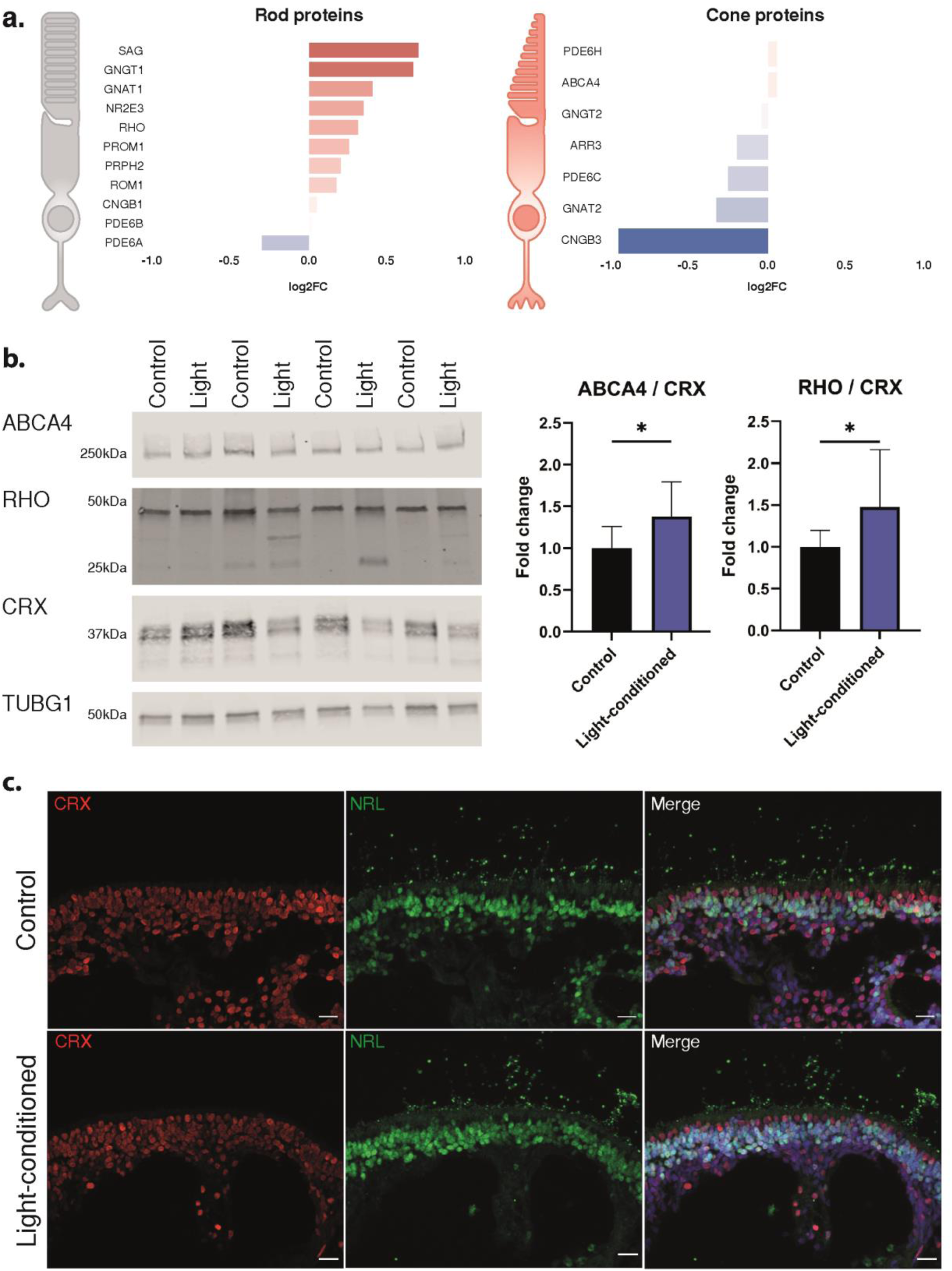
Rod photoreceptor protein markers are enriched in light-conditioned ROs. **(a)** Representation of the log2 fold change of rod (left) and cone (right) protein markers, detected in the performed proteomics screen, showing enrichment of rod markers. **(b)** Western blot analysis of several photoreceptor markers, accompanied by a semi-quantification (right panel) after normalization against CRX to correct for photoreceptor content. The mean ± SD of four samples with three technical replicates are displayed. **(c)** Immunofluorescent analysis of DIV210 control and light-conditioned ROs for photoreceptor transcription factors CRX (red) and NRL (green), marking cones (CRX+, NRL-) and rods (CRX+, NRL+), respectively. Cell nuclei were stained using DAPI (blue). Scale bar = 20 µm.

### Long-term light conditioning does not lead to increased stress in ROs

Finally, we evaluated whether light conditioning caused stress or cell death in ROs. Light damage is known to cause increased expression of apoptotic, autophagy, and stress markers.^24^ We therefore analysed the expression of these markers in our proteomics dataset. From our dataset, we were only able to include proteins that are known to increase in expression, as we cannot distinguish between inactive or activated variants of a protein. Based on this, we created a list of five markers for each cellular process, i.e., apoptosis (AIP, BAK, BAX, BNIP3, FAS), autophagy (ATG3, ATG5, ATG7, BECN1, LAMP2), and stress (BIP, GRP94, HMOX1, SOD1, SOD2), respectively. Light conditioning did not result in changes in the expression of pro-apoptotic (**Figure 5a**), autophagy (**Figure 5b**), and stress markers (**Figure 5c**). We followed up on these findings by measuring the expression of ER stress markers *BIP* and *GRP94* by qPCR to assess whether light conditioning results in increased stress. As a positive control for our readout, we treated DIV209 control ROs for 48 hours with 2 µg/mL Tunicamycin (TM), a known inhibitor of glycosylation that leads to increased ER stress.^25^ As expected, TM treatment led to an approximately 10-fold increase in gene expression of stress markers using qPCR, showing the capacity of ROs to generate stress responses (**Figure 5d**). In contrast, we did not observe increased expression of stress related genes *BIP* and *GRP94* in our light-conditioned ROs even after 20 weeks (DIV70-DIV210) of light conditioning. Lastly, to further assess apoptosis in light-conditioned ROs, we stained DIV210 ROs for active Caspase-3 (CASP3). Here, we did not observe noticeable changes in CASP3 expression around CRX+ photoreceptor nuclei, while CASP3+ cells were present in TM-treated ROs (**Figure 5e**), showing that light conditioning did not result in light damage or stress in ROs using our setup.

**Figure 5.**
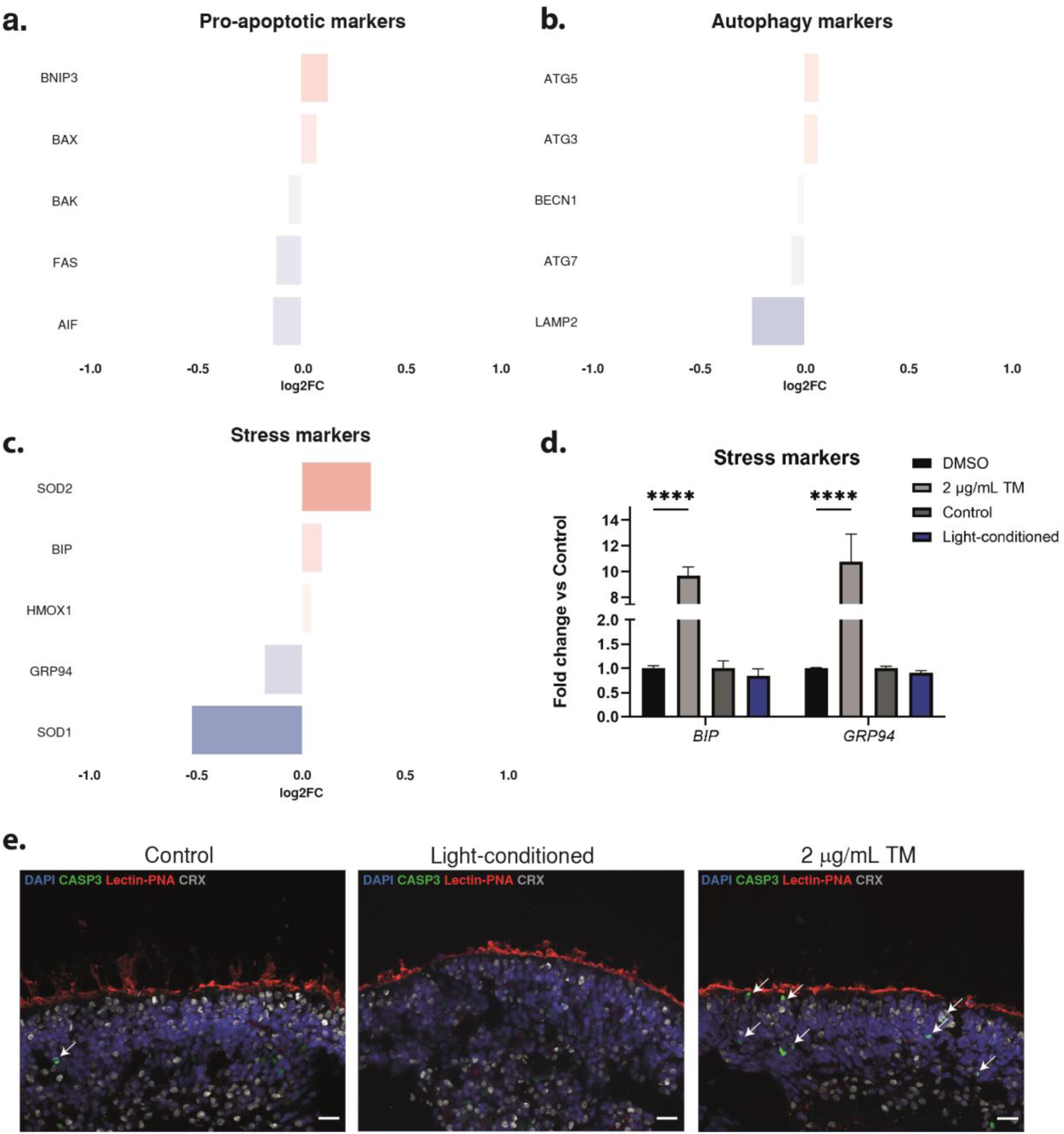
Light-conditioned ROs do not show increased stress or cell death compared to control. **(a)** Analysis of pro-apoptotic protein markers based on the proteomic dataset between control and light-conditioned ROs. **(b)** Analysis of autophagy protein markers based on the proteomic dataset between control and light-conditioned ROs. **(c)** Analysis of stress proteins markers based on the proteomic dataset between control and light-conditioned ROs. **(d)** Assessment of stress markers (*BIP* and *GRP94*) by qPCR in DIV210 control and light-conditioned ROs in two independent RO differentiation batches with three technical replicates each. As positive and negative control, ROs treated with 2 µg/ml tunicamycin (TM) or DMSO were used, respectively. TM treatment was performed in one differentiation batch using three technical replicates. Bars represent the fold change mean ± SD against the DMSO control. Statistical differences are indicated with asterisks (****p-value<0.0001) using one-way ANOVA followed by a post-hoc test. **(C)** Immunofluorescence analysis of DIV210 control and light-conditioned ROs for cell death marker cleaved Caspase-3 (CASP3) (green), Lectin-PNA (red), CRX (grey), and DAPI (blue). White arrows indicate CASP3+ cells. Scale bar = 20 µm.

## Discussion

In this study, we employed long-term white light conditioning to enhance the maturation of photoreceptors in human iPSC-derived ROs. We showed that six hours of daily light accelerated the expression of photoreceptor gene levels over time, as well as increasing photoreceptor OS length, indicating accelerated photoreceptor generation compared to control. Furthermore, light-conditioned ROs showed increased (rod) photoreceptor gene and protein expression levels at DIV210, without increasing stress and cell death. Our results show that light conditioning enhances photoreceptor maturation in ROs.

The role of light in retinal development has been previously investigated in animal models, where the lack of light led to disrupted photoresponses.^19–21^ General RO differentiation protocols are mainly performed in a dark incubator and therefore could lead to disrupted or delayed retinal development. By supplementing light in RO culturing, we were able to enhance photoreceptor maturation. These changes are likely to be driven by the expression of transcription factors important for the development of the retina. Although we show that *CRX* gene expression is enhanced at DIV210, protein levels remained stable in our proteomics dataset. Interestingly, researchers have found that CRX expression is post-transcriptionally regulated by light in the developing rat retina, but not in the adult retina.^26^ This could indicate that light indeed influences CRX expression, and consequently other downstream photoreceptor markers essential for proper retinal development.

Interestingly, the role of light in human iPSC-derived ROs has not been extensively studied. However, it was previously demonstrated that light, and also different wavelengths of light, regulate the expression of miRNAs, including the miRNA cluster miR-182/183/96 associated with retinal development.^27^ Additionally, complementing and supporting our findings, Celiker *et al.* (submitted to BioRxiv on the same day as this work) reported that light stimulation not only enhances photoreceptor differentiation in ROs, but that light also changes the cellular composition towards a more mature neuronal profile after seven days of 12-hour cycles of dark and light using a 40 Hz flickering light. While researchers successfully showed changes in this setting, we observed that long-term continuous light in a 12-hour dark/light cycle does not improve RO differentiation, but rather that six hours of daily light appears to be sufficient to accelerate photoreceptor generation in ROs. These discrepancies can be caused by several differences in the respective experimental set-up, including the way of light stimulation (bottom versus top), the type of light (flickering versus constant), RO differentiation protocol, media type (without phenol red versus with phenol red), the starting timepoint (mature versus DIV70), as well as the duration of the exposure (one week versus 20 weeks). However, both studies converge on the idea that light enhances the maturation of photoreceptors in ROs, highlighting the crucial role of light in retinal development and maturation.

Light-conditioned ROs specifically exhibited an increased proportion of rod photoreceptors, as indicated by increased rod-specific protein levels. We also observed relatively decreased protein levels of cone-specific markers, however these changes could also be explained by an increased ratio of rod:cone photoreceptor cells, which could indicate a more advanced developmental stage, more similar to the totally formed retina for which a rod:cone ratio of 20:1 is generally observed.^28,29^ Another explanation could be due to the known retinal developmental temporal order. Rod photoreceptors cells are one of the last cell types to develop in the mammalian retina. While in mice, cone photoreceptors arise in embryonic stages, and rod photoreceptors are mainly formed postnatally, together with bipolar cells and Müller glia.^30^ By DIV65, the pool of proliferating *KI67*+ cells is halved in ROs, and cone photoreceptor markers are appearing.^31^ Therefore, the remaining retinal progenitor cells may be limited to differentiate into the late-born rod photoreceptors. Future studies should include proteomic analysis at different time-points to detect whether indeed cones also benefit from the light stimulation, but the effect is buffered by the fact that the population of rod photoreceptors increases.

Despite the observed significant positive changes in photoreceptor differentiation in light-conditioned ROs (based on OS measurements and qPCR data), we found limited changes on protein level using proteomics. These changes are likely due to a) a lack of linearity between RNA expression and protein synthesis; and b) the known heterogeneity between ROs and RO differentiation in the different batches. We aimed to correct these issues by pooling multiple sets of ROs to get an accurate representation of the tested conditions. Doing so, we were able assess differences in the expression of specific markers and observed a clear increase in the expression of rod markers in the proteomics data.

One of the big challenges on the use of ROs is the time-consuming and long differentiation protocols which go beyond DIV210, together with the lack of disease phenotypes compared to human conditions (e.g. lack of photoreceptor degeneration). Our findings that light conditioning increases photoreceptor maturation in ROs could be important for disease modelling and therapy development, especially for IRDs. Specifically, many of the IRD genes are expressed in photoreceptor OS and are important in light processing.^32–34^ As a result, implementing light exposure in the differentiation of ROs harboring pathogenic variants will likely facilitate the detection of molecular and cellular defects specifically related to the function of the respective gene.

Our research shows that light conditioning using constant white light (350-400 LUX) improves the expression of photoreceptors in ROs. Future directions could go into deciphering which light wavelength(s) have a more prominent effect on RO differentiation, and whether changes in the light intensity could increase or decrease the maturation of ROs. Still, here we present the first steps towards understanding how light can improve the use of ROs for disease modeling and therapeutic intervention. Increasing photoreceptor maturation may allow for the development of more robust functional readouts. Previous studies have shown that cone photoreceptors in ROs are able to exhibit light-evoked responses, similarly to the macaque cone photoreceptors.^35^ However, these responses remain limited, likely due to underdeveloped photoreceptor OSs and the immature nature of ROs.^14,15^ Therefore, functionally assessing whether light-responses are improved in light-conditioned ROs using for example patch-clamp of individual photoreceptors, will be of great added value.

In summary, exposing human iPSC-derived ROs to light on a daily basis improved the generation and maturation of photoreceptors, without detrimental effects. Therefore, implementation of daily light in differentiation protocols can provide more relevant IRD models and thereby accelerate therapy development.

## Methods

### iPSC culturing and retinal organoid differentiation

Control iPSCs were characterized previously and were cultured in Essential 8 Flex (E8F) media (Gibco, A2858501) on Geltrex (Gibco, A1413302)-coated plates at 37 °C and 5% CO2.^36^ For all experiments, the SCTCi008-A iPSC line was used. iPSCs were passed every 4-7 days for maintenance. Differentiation of iPSCs into ROs was based on a previously published protocol, with some adjustments^37^. Briefly, iPSCs were detached from the culture plates in large clumps and were allowed to form embryoid bodies overnight in a T25 flask. The following days, the Essential 8 Flex media was gradually replaced with neural induction media (NIM). At DIV6, 1.5 nM BMP4 (Sigma, SRP3016) was added to NIM, followed by half media changes on DIV8, 10 and 13. On DIV8, EBs were plated in a 6-well plate which was pre-treated with fetal calf serum (FCS) to ensure attachment. On DIV16, attached cells were scraped off and placed in retinal differentiation media (RDM) in an ultra-low attachment plate (Corning, 3471). At DIV36, media was replaced with retinal maturation media (RMM). Different supplements were added to RMM at different timepoints; 1x B27 supplement (Gibco, 12587010) (DIV36 - DIV100), 1 µM Retinoic Acid (DIV50 – DIV70), 0.5 µM Retinoic Acid (DIV71 – DIV100), 1x N2 supplement (Gibco, 17502048) (DIV71 – end of culture). A full media composition list can be found in **Supplementary Table 1**. An overview of the RO differentiation protocol can be found in **Figure 1a**.

### Light conditioning of retinal organoids

Full spectrum LED lights were lined along one of the shelves within the incubator, in order for the lights to shine from the top (**Figure 1b**). Light intensity was measured using a LUX meter and was between 350-400 LUX across the shelf. Control ROs were cultured on the top shelf of the same incubator, where light intensity was measured at 0 LUX, even with the lights on the lower shelf turned on. ROs were exposed to light based on an automatic timer every day for 6 hours for most of the experiments (**Figure 1c**). Only for the pilot test to see how much light the ROs can handle, 12 hours of daily light was used. Light conditioning was performed from DIV70 onwards. For every experiment, ROs (both light-conditioned and control) were harvested in the middle of the light cycle of the light-conditioned ROs.

### RNA isolation, cDNA synthesis, and qPCR

For RNA experiments, five ROs were pooled during harvesting. Samples were snap frozen in liquid nitrogen after washing the retinal organoids with PBS, and stored at −80 °C until use. RNA was isolated using the NucleoSpin RNA kit (Macherey-Nagel, MN740955), according to manufacturer’s instructions. After RNA isolation, RNA was precipitated by adding 0.5 volumes of 5M ammonium acetate (Invitrogen, AM9070G), 2.5 volumes of ice-cold 100% ethanol, and 1 µL Glycogen (Invitrogen, 10814010), followed by overnight incubation at −20 °C. The following day, RNA was recovered by centrifugation at 4 °C for 30 minutes at 12.000 x *g*. The supernatant was removed, and the RNA was washed by adding 0.5 mL ice-cold 70% ethanol, and centrifuging at 4 °C for 10 minutes at 12.000 x *g*. The supernatant was removed again, and the tubes were left open at room temperature until the last traces of ethanol evaporated. Finally, the RNA was resuspended in nuclease-free water to the desired volume. cDNA synthesis was performed using the SuperScript VI (Invitrogen, 11755250), according to manufacturer’s instructions. A maximum of 1000 ng RNA was used for one reaction. cDNA was then diluted to 5 ng/µL for qPCR analysis. The qPCR reaction was set-up using the GoTaq qPCR master mix (Promega, A6002), according to the manufacturer’s instructions. The final cDNA concentration was 1 ng/µL per reaction. Samples were run on a QuantStudio 5 Real-Time PCR system. Raw CT values were normalized against the housekeeping gene *GUSB*. A full list of all the target genes and corresponding primer sequences is listed in **Supplementary Table 2**.

### Protein isolation

For both the western blot and proteomic experiments, the same protocol for protein isolation was used. Three ROs were pooled per condition. Samples were briefly washed with PBS, before snap-freezing in liquid nitrogen and stored at −80 °C until use. ROs were lysed in RIPA buffer, containing 0.75% SDS. ROs were disrupted by addition of a glass bead followed by shaking withing the TissueLyser II (Qiagen) at a frequency of 1800 oscillations per minute for 1 minute. Samples were then incubated on ice for 1 hour, before sonification of the samples. Next, samples were spun down at max speed for 10 minutes at 4 °C, and supernatant was moved to a new tube. Protein concentration was determined using the Pierce BCA protein assay kit (Thermo Scientific, 23225), according to manufacturer’s instructions.

### Proteomics

Extracts containing 10 µg protein were diluted 1:1 with 8 M urea 10 mM Tris-HCl (pH8) prior to detergent removal using Pierce detergent removal spin columns (Thermo Fisher scientific) according to manufacturer’s recommendations. Reduction and alkylation of cysteine residues was done by addition of 1 µl 50 mM dithiothreitol and 1 µl 50 mM chloroacetamide and subsequent incubation at room temperature for 30 minutes in the dark, respectively. Next, 50 µl 50 mM ammonium bicarbonate (pH 8) and 0.5 µl Trypsin (0.2 µg/µl) were added and samples were incubated overnight at 37 °C. Tryptic digests were analyzed by nanoflow liquid chromatography (Evosep One, Evosep Biosystems) coupled online to a trapped ion mobility spectrometry – quadrupole time-of-flight mass spectrometer (timsTOF Pro2, Bruker Daltonics) via a nanoflow electrospray ionization source (CaptiveSprayer, Bruker Daltonics). Tryptic peptides were separated by C18 reversed phase liquid chromatography (Evosep EV1137 30SPD performance column; 150 mm length x 0.150 mm internal diameter, 1.5 µm C18AQ particles) using the pre-programmed 30 samples per day (30SPD) Evosep One method. The mass spectrometer was operated in positive ionization mode using the default data independent acquisition – Parallel Accumulation SErial Fragmentation (dia-PASEF) ^38^ instrument method: 0.6 – 1.6 1/K0 mobility range, 100 - 1700 m/z mass range, 100 ms accumulation time, 100 ms ramp time, 26 Da mass width, 1 Da mass overlap, 32 mass steps per cycle, 0 mobility overlap, 1 mobility window. Acquired spectra were streamed directly to ProteoScape (v2025b, Bruker Daltonics) for protein identification and label-free quantitation against the Uniprot Homo sapiens canonical protein sequence database (downloaded Jan 2024) using the following settings: Spectronaut v19 directDIA+ (Fast) workflow, 0.2 precursor PEP cutoff, 0.01 precursor Q-value cutoff, 0.01 protein Q-value cutoff global, 0.01 protein Q-value cutoff, 0.75 protein PEP cutoff, full tryptic specificity, allowed up to 2 missed cleavages, carbamidomethyl (C) as fixed modification and Oxidation (M) as variable modifications, protein group specific peptides were used for quantitation. Of all detected proteins, we filtered out the proteins that were only detected in two or less samples, leaving only the remaining reliably detected proteins. The log2 fold change was calculated for the light-conditioned samples compared to the control samples. A principal component analysis of all detected proteins was generated. A heatmap was generated using the differentially expressed proteins. A volcano plot was generated where proteins with a p-value < 0.05 and log2FC ≥ 0.5 were marked as significant. Data analysis and visualization were performed in R version 4.4.1. using the following packages: tidyverse version 2.0.0, ggplot2 version 3.5.2, pheatmap version 1.0.13, FactoMineR version 2.11, factoextra version 1.0.7, dplyr version 1.1.4, DESeq2 version 1.46.0, and missMDA version 1.18. Gene Ontology analysis was performed using the online ShinyGO V0.82 tool.^39^ Here, all detected proteins in our dataset were used as a background and a false discovery rate of 0.05 was used to find disrupted pathways.

### SDS-PAGE and Western blotting

A total of 20 µg of protein was supplemented with 4x Laemmli sample buffer (Bio-Rad, 1610747) and 10% DTT before loading on a 4-15% Mini-PROTEAN TGX Stain-Free Gel (Bio-Rad, 4568084). Samples were run for 30 minutes at 250 V in 1x TRIS/SDS/Glycine running buffer. Proteins were transferred to a nitrocellulose membrane (Bio-Rad, 1704158) using a mixed molecular weight program on the Trans-Blot Turbo transfer system (Bio-Rad, 1704150). The nitrocellulose membranes were blocked for 1 hour in 5% non-fat dry milk in PBS. Blots were incubated with primary antibodies diluted in 5% non-fat dry milk overnight. Membranes were then washed three times using PBS+ 0.2% Tween 20 before incubation with secondary antibodies for 2 hours in 5% non-fat dry milk in PBS. A list of all primary and secondary antibodies used, including concentrations, can be found in **Supplementary Table 3**. Blots were scanned using the Odyssey CLx Imager (LI-COR).

### Organoid fixation and immunofluorescence

At least three ROs were collected per condition for immunofluorescent analysis. ROs were washed twice in PBS before fixation in 4% PFA for 20 minutes at room temperature. Following PBS washes, ROs were suspended in 30% glucose in PBS solution at 4 °C until the ROs were saturated and settled at the bottom, approximately two hours, for cryopreservation. ROs were then embedded in Tissue-Tek O.C.T. compound and frozen on dry ice. Next, 12 µm thick sections were made using a cryostat, and slides were frozen at −20 °C until use. For immunofluorescent labeling, retinal organoid sections were rehydrated in PBS for 20 minutes, before permeabilization with 0.1% Triton X-100 in PBS for 15 minutes. After permeabilization, sections were blocked in blocking buffer containing 10% Normal Donkey Serum, 1% Bovine Serum Albumin in 0.3% Triton X-100 in PBS for 1 hour at room temperature. Primary antibodies were diluted in blocking buffer, and added to the slides to incubate overnight at 4 °C. After three 10-minute PBS washes, secondary antibodies, diluted in PBS, were added to the slides for two hours at room temperature, to allow for fluorescent detection. RO nuclei were labeled using DAPI. Images were taken with the Zeiss Axio Imager, equipped with an apotome, and images were processed using FIJI 1.53t. A full list of primary and secondary antibodies used is provided in **Supplementary Table 3**.

### Statistical analysis

For gene expression data of DIV105 ROs, four independent differentiation batches of retinal organoids with three technical replicates each were used. Significance was calculated using an unpaired t-test, to compare control and light-conditioned ROs. For gene expression data up to DIV210, two independent batches of ROs were used, with three technical replicates each. A two-way ANOVA was followed by a post-hoc test to determine differences between control and light-conditioned ROs and between timepoints. Differences in the size of the POS were determined using a Mann-Whitney test, as data was not normally distributed. For proteomics data, a total of four pools of ROs were submitted per condition, and each sample was ran three times. From the three technical replicates, the averages were calculated and used for further significance testing as described in the proteomics section. Statistical analysis for all data except proteomics data was done using GraphPad Prism 10. All results with a p-value of < 0.05 were considered significant.

## Data availability

The mass spectrometry proteomics data have been deposited to the ProteomeXchange Consortium via the PRIDE ^40^ partner repository with the dataset identifier PXD065849.

## Acknowledgements.

We would like to thank the Collin-Garanto lab members for feedback and sporadic help with experimental work. We would like to thank Dirk Lefeber and Susan Roelofs for discussions and feedback on scientific content, as well as all the members of the RoC-ME consortium. We would like to also thank Fokje Zijlstra and Hans Wessels for their help with the sample preparation and analysis of the proteomics data. This work was financed by the Human Measurement Models call 2.0 (grant nr. 18958 (RoC-ME) to AG and AvdM) with additional funding from Proefdiervrij; and it is supported by the Association of Collaborating Health Foundations (SGF), NWO Domain AES and the Netherlands Organisation for Health Research and Development (ZonMw), as part of their joint strategic research programme: Human Measurement Models”. The collaboration project is co-funded by the PPP Allowance made available by Health∼Holland, Top Sector Life Sciences & Health, to the Association of Collaborating Health Foundations (SGF) to stimulate public-private partnerships. Additional funding has been obtained through contributions from health foundations and supported in part by interested industrial research partners. Proteomics measurements were performed by the Radboud Technology Center for Mass Spectrometry supported by the Netherlands X-omics Initiative partially funded by NWO (project 184.034.019). The iPSC cells were generated by the Radboudumc Stem Cell Technology Center (https://www.radboudumc.nl/en/research/radboud-technology-centers/stem-cells).

## Author contributions

Conceptualization: E.v.O., R.C., and A.G. Methodology: E.v.O., A.H., W.K., S.B., J.H., M.v.d.M., and A.G. Scientific discussions: E.v.O., A.H., W.K., S.B., J.H., M.v.d.M., D.V., A.B., S.A., A.v.d.M., T.B., R.C., and A.G. Investigation: E.v.O., A.H., W.K., S.B., J.H., M.v.d.M., and A.G. Funding acquisition: S.A., A.v.d.M., and A.G. Supervision: R.C., A.G. Writing original draft: E.v.O., A.G., Review and editing: E.v.O, A.H., W.K., S.B., J.H., M.v.d.M., D.V., A.B., S.A., A.v.d.M., T.B., R.C. and A.G.

## References

1 Dowden, H. & Munro, J. Trends in clinical success rates and therapeutic focus. Nat Rev Drug Discov 18, 495–496 (2019). 10.1038/d41573-019-00074-z

2 Nadal-Nicolas, F. M. et al. True S-cones are concentrated in the ventral mouse retina and wired for color detection in the upper visual field. Elife 9 (2020). 10.7554/eLife.56840

3 Langmann, T. et al. Comparison of Mouse and Human Retinal Pigment Epithelium Gene Expression Profiles: Potential Implications for Age-Related Macular Degeneration. Plos One 10 (2015). 10.1371/journal.pone.0141597

4 Liu, X. et al. Usherin is required for maintenance of retinal photoreceptors and normal development of cochlear hair cells. Proc Natl Acad Sci U S A 104, 4413–4418 (2007). 10.1073/pnas.0610950104

5 Charbel Issa, P., et al. Fundus autofluorescence in the Abca4(-/-) mouse model of Stargardt disease--correlation with accumulation of A2E, retinal function, and histology. Invest Ophthalmol Vis Sci 54, 5602–5612 (2013). 10.1167/iovs.13-11688

6 Bharti, K. et al. Cell culture models to study retinal pigment epithelium-related pathogenesis in age-related macular degeneration. Exp Eye Res 222, 109170 (2022). 10.1016/j.exer.2022.109170

7 Zhu, Y. et al. In vitro Model Systems for Studies Into Retinal Neuroprotection. Front Neurosci 16, 938089 (2022). 10.3389/fnins.2022.938089

8 Rowe, R. G. & Daley, G. Q. Induced pluripotent stem cells in disease modelling and drug discovery. Nature Reviews Genetics 20, 377–388 (2019). 10.1038/s41576-019-0100-z

9 Kaiser, V. M. & Gonzalez-Cordero, A. Organoids - the future of pre-clinical development of AAV gene therapy for CNS disorders. Gene Ther (2025). 10.1038/s41434-025-00527-8

10 Afanasyeva, T. A. V. et al. A look into retinal organoids: methods, analytical techniques, and applications. Cellular and Molecular Life Sciences 78, 6505–6532 (2021). 10.1007/s00018-021-03917-4

11 Zhang, X., Wang, W. & Jin, Z. B. Retinal organoids as models for development and diseases. Cell Regen 10, 33 (2021). 10.1186/s13619-021-00097-1

12 Chakrabarty, K. et al. Retinal organoids in disease modeling and drug discovery: Opportunities and challenges. Surv Ophthalmol 69, 179–189 (2024). 10.1016/j.survophthal.2023.09.003

13 Zhao, H. & Yan, F. Retinal Organoids: A Next-Generation Platform for High-Throughput Drug Discovery. Stem Cell Reviews and Reports 20, 495–508 (2023). 10.1007/s12015-023-10661-8

14 Sridhar, A. et al. Single-Cell Transcriptomic Comparison of Human Fetal Retina, hPSC-Derived Retinal Organoids, and Long-Term Retinal Cultures. Cell Rep 30, 1644–1659 e1644 (2020). 10.1016/j.celrep.2020.01.007

15 Wahle, P. et al. Multimodal spatiotemporal phenotyping of human retinal organoid development. Nat Biotechnol 41, 1765–1775 (2023). 10.1038/s41587-023-01747-2

16 Georgiou, M., Fujinami, K. & Michaelides, M. Inherited retinal diseases: Therapeutics, clinical trials and end points-A review. Clin Exp Ophthalmol 49, 270–288 (2021). 10.1111/ceo.13917

17 Gong, J. et al. A controllable perfusion microfluidic chip for facilitating the development of retinal ganglion cells in human retinal organoids. Lab Chip 23, 3820–3836 (2023). 10.1039/d3lc00054k

18 Inagaki, S. et al. Establishment of vascularized human retinal organoids from induced pluripotent stem cells. Stem Cells 43 (2025). 10.1093/stmcls/sxae093

19 Akimov, N. P. & Renteria, R. C. Dark rearing alters the normal development of spatiotemporal response properties but not of contrast detection threshold in mouse retinal ganglion cells. Dev Neurobiol 74, 692–706 (2014). 10.1002/dneu.22164

20 Bonezzi, P. J., Tarchick, M. J., Moore, B. D. & Renna, J. M. Light drives the developmental progression of outer retinal function. J Gen Physiol 155 (2023). 10.1085/jgp.202213262

21 Chai, Z. et al. Light-dependent photoreceptor orientation in mouse retina. Sci Adv 6 (2020). 10.1126/sciadv.abe2782

22 Hofmann, K. P. & Lamb, T. D. Rhodopsin, light-sensor of vision. Prog Retin Eye Res 93, 101116 (2023). 10.1016/j.preteyeres.2022.101116

23 Klapper, S. D., Swiersy, A., Bamberg, E. & Busskamp, V. Biophysical Properties of Optogenetic Tools and Their Application for Vision Restoration Approaches. Front Syst Neurosci 10, 74 (2016). 10.3389/fnsys.2016.00074

24 Ouyang, X. et al. Mechanisms of blue light-induced eye hazard and protective measures: a review. Biomed Pharmacother 130, 110577 (2020). 10.1016/j.biopha.2020.110577

25 Wang, S. et al. Tunicamycin-induced photoreceptor atrophy precedes degeneration of retinal capillaries with minimal effects on retinal ganglion and pigment epithelium cells. Exp Eye Res 187, 107756 (2019). 10.1016/j.exer.2019.107756

26 Wu, Y. et al. Crx Is Posttranscriptionally Regulated by Light Stimulation in Postnatal Rat Retina. Front Cell Dev Biol 8, 174 (2020). 10.3389/fcell.2020.00174

27 Celiker, C. et al. Light-responsive microRNA molecules in human retinal organoids are differentially regulated by distinct wavelengths of light. iScience 26, 107237 (2023). 10.1016/j.isci.2023.107237

28 Curcio, C. A., Sloan, K. R., Kalina, R. E. & Hendrickson, A. E. Human photoreceptor topography. J Comp Neurol 292, 497–523 (1990). 10.1002/cne.902920402

29 Hussey, K. A., Hadyniak, S. E. & Johnston, R. J., Jr. Patterning and Development of Photoreceptors in the Human Retina. Front Cell Dev Biol 10, 878350 (2022). 10.3389/fcell.2022.878350

30 Bassett, E. A. & Wallace, V. A. Cell fate determination in the vertebrate retina. Trends Neurosci 35, 565–573 (2012). 10.1016/j.tins.2012.05.004

31 Agarwal, D. et al. Bulk RNA sequencing analysis of developing human induced pluripotent cell-derived retinal organoids. Sci Data 9, 759 (2022). 10.1038/s41597-022-01853-x

32 Hu, H. et al. Cross-species single-cell landscapes identify the pathogenic gene characteristics of inherited retinal diseases. Front Genet 15, 1409016 (2024). 10.3389/fgene.2024.1409016

33 Comitato, A., Schiroli, D., La Marca, C. & Marigo, V. Differential Contribution of Calcium-Activated Proteases and ER-Stress in Three Mouse Models of Retinitis Pigmentosa Expressing P23H Mutant RHO. Adv Exp Med Biol 1185, 311–316 (2019). 10.1007/978-3-030-27378-1_51

34 Molday, R. S., Garces, F. A., Scortecci, J. F. & Molday, L. L. Structure and function of ABCA4 and its role in the visual cycle and Stargardt macular degeneration. Prog Retin Eye Res 89, 101036 (2022). 10.1016/j.preteyeres.2021.101036

35 Saha, A. et al. Cone photoreceptors in human stem cell-derived retinal organoids demonstrate intrinsic light responses that mimic those of primate fovea. Cell Stem Cell 29, 460–471 e463 (2022). 10.1016/j.stem.2022.01.002

36 Koolen, L. et al. Generation and characterization of human induced pluripotent stem cells (iPSCs) from three individuals without age-related macular degeneration. Stem Cell Res 60, 102670 (2022). 10.1016/j.scr.2022.102670

37 Fligor, C. M., Huang, K. C., Lavekar, S. S., VanderWall, K. B. & Meyer, J. S. Differentiation of retinal organoids from human pluripotent stem cells. Methods Cell Biol 159, 279–302 (2020). 10.1016/bs.mcb.2020.02.005

38 Meier, F. et al. diaPASEF: parallel accumulation-serial fragmentation combined with data-independent acquisition. Nat Methods 17, 1229–1236 (2020). 10.1038/s41592-020-00998-0

39 Ge, S. X., Jung, D. & Yao, R. ShinyGO: a graphical gene-set enrichment tool for animals and plants. Bioinformatics 36, 2628–2629 (2020). 10.1093/bioinformatics/btz931

40 Perez-Riverol, Y. et al. The PRIDE database at 20 years: 2025 update. Nucleic Acids Res 53, D543–D553 (2025). 10.1093/nar/gkae1011

