## Supplementary Material for "Long-term daily light exposure boosts photoreceptor maturation in retinal organoids"

[1.] Radboud university medical center, Amalia Children's Hospital, Department of Pediatrics, Nijmegen, The Netherlands
[2.] Radboud university medical center, Department of Human Genetics, Nijmegen, The Netherlands
[3.] Applied Stem Cell Technologies Group, Department of Bioengineering Technologies, University of Twente, Enschede, The Netherlands
[4.] BIOS Lab on a Chip group, MESA+ Institute for Nanotechnology, University of Twente, Enschede, The Netherlands
[5.] Eye Health & Research Beyond Borders, Boehringer Ingelheim Pharma GmbH & Co. KG, Biberach, Germany
[6.] Department of Histology and Embryology, Faculty of Medicine, Masaryk University, Brno, Czech Republic

[*] Corresponding author

To whom correspondence should be addressed:
Dr. Alejandro Garanto
Dept. Pediatrics and Dept. Human Genetics (route 855)
Geert Grooteplein 10
6525 GA Nijmegen, The Netherlands


**Supplementary Table 1.** Overview of retinal organoid differentiation media.

| Media type | Base media | Supplements |
|---|---|---|
| NIM | Advanced DMEM/F12 (Gibco, 12634010) | 1x N2 supplement (Gibco, 17502048), 1x Glutamax (Gibco, 35050038), 2µg/mL Heparin (Sigma, H3149), 100µg/mL Primocin (Invitrogen, ant-pm-2) |
| RDM | DMEM (Gibco, 41966052) / F-12 Ham (Sigma-Aldrich, N6658) (3:1 ratio) | 1x B27 supplement (Gibco, 12587010), 1x MEM-Non Essential Amino Acids (Sigma, M7145), 100µg/mL Primocin (Invitrogen, ant-pm-2) |
| RMM | DMEM (Gibco, 41966052) / F-12 Ham (Sigma-Aldrich, N6658) (3:1 ratio) | 10% FCS, 1x Glutamax (Gibco, 35050038), 100µM Taurine (Sigma, T0625), 100µg/mL Primocin (Invitrogen, ant-pm-2) |

**Supplementary Table 2.** qPCR primer sequences used in this study.

| Primer name | Sequences (5'-3') |
|---|---|
| *ARR3* F | GTGGAACCCATTGACGGTG |
| *ARR3* R | CACTTCCAAGTCATCACGGC |
| *BIP* F | AGAACCAGCTCACCTCCAAC |
| *BIP* R | GTCTTTGTTTGCCCACCTCC |
| *CRX* F | CCCCAGTGTGGATCTGATG |
| *CRX* R | CAAACAGTGCCTCCAGCTC |
| *GRP94* F | GGAGAGTCGTGAAGCAGTTG |
| *GRP94* R | TGTAGAGATGTCCTTGCCCG |
| *GUSB* F | AGAGTGGTGCTGAGGATTGG |
| *GUSB* R | CCCTCATGCTCTAGCGTGTC |
| *PDE6H* F | TCCCAAGTTCAAGCAGAGG |
| *PDE6H* R | GTTCCTAGCCCCTCCATTC |
| *PRPH2* F | CTCTTGATTCGATCTTTGAC |
| *PRPH2* R | ACATGCTGCAGATCGAGTTC |
| *RCVRN* F | ACACCAAGTTCTCGGAGGAG |
| *RCVRN* R | ACTTGGCGTAGATGCTCTGG |
| *RHO* F | TCATCATGGTCATCGCTTTC |
| *RHO* R | CATGAAGATGGGACCGAAGT |

**Supplementary Table 3.** Antibodies used in this study.

| Target | Antibody | Dilution per application |
|---|---|---|
| ABCA4 | Rabbit Anti-ABCA4 (Abcam, ab72955) | WB: 1:1.000 |
| CRX | Mouse Anti-CRX (Abnova, H00001406-M02) | IF: 1:500, WB: 1:2.000 |
| Active Caspase-3 | Rabbit Anti-Active Caspase-3 ( BD Pharmingen, 559565) | IF: 1:500 |
| Lectin-PNA | Lectin-PNA, Alexa Fluor 568 Conjugate (Invitrogen, L32458) | IF: 1:100 |
| NRL | Goat Anti-NRL (R&D Systems, AF2945) | IF: 1:500 |
| RHO | Mouse Anti-Rhodopsin (4D2) (Novus Biologicals, NBP2-59690) | WB: 1:5.000 |
| Gamma-Tubulin | Mouse Anti-Gamma Tubulin (Sigma-Aldrich, T5326) | WB: 1:2.000 |
| Secondary | Goat Anti-Rabbit Alexa Fluor 680 (Invitrogen, A21076) | WB: 1:10.000 |
| Secondary | Goat Anti-Mouse IRDye 800CW (Licor Biosciences, 926-32210) | WB: 1:10.000 |
| Secondary | Goat Anti-Mouse Alexa Fluor 680 (Invitrogen. A21057) | WB: 1:10.000 |
| Secondary | Goat Anti-Rabbit IRDye 800CW (Licor Biosciences, 926-32211) | WB: 1:10.000 |
| Secondary | Donkey Anti-Rabbit Alexa Fluor 488 (Invitrogen, A21206) | IF: 1:500 |
| Secondary | Donkey Anti-Mouse Alexa Fluor 647 (Invitrogen, A31571) | IF: 1:500 |
| Secondary | Donkey Anti-Goat Alexa Fluor 488 (Invitrogen, A11055) | IF: 1:500 |
| Secondary | Donkey Anti-Mouse Alexa Fluor 568 (Invitrogen, A10037) | IF: 1:500 |
| Secondary | Donkey Anti-Rabbit Alexa Fluor 647 (Invitrogen, A31573) | IF: 1:500 |
| DAPI | DAPI (Invitrogen, D1306) | IF: 1:5000 |

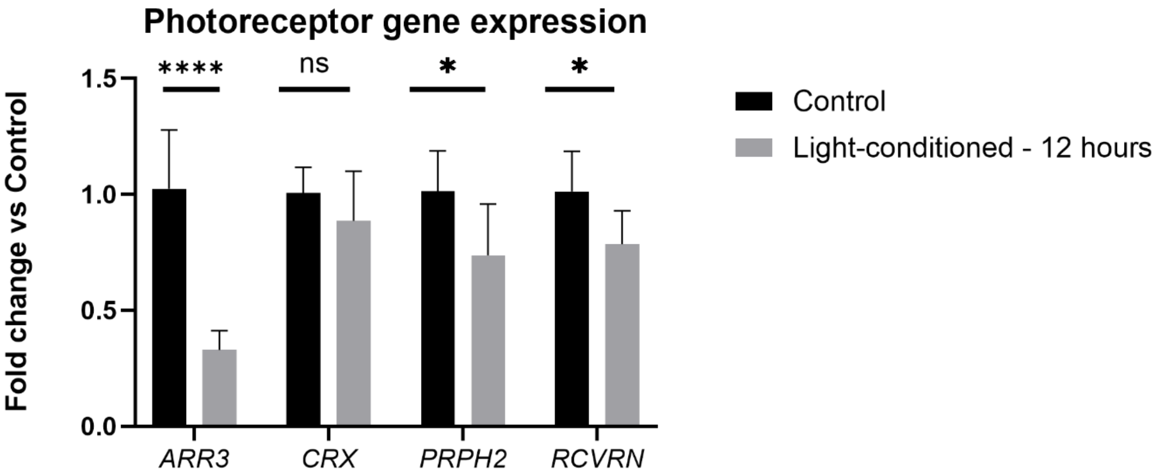


**Supplementary Figure 1. qPCR of photoreceptor genes after 5-week of 12-hour light conditioning.** Gene expression analysis of photoreceptor markers from one RO differentiation with three technical replicates each. Light-conditioned organoids showed a significant decrease compared to control ROs. Values are represented as the mean ± SD. Statistical differences are indicated with asterisk (* p-value<0.05; **** p-value<0.0001) using an unpaired t-test.

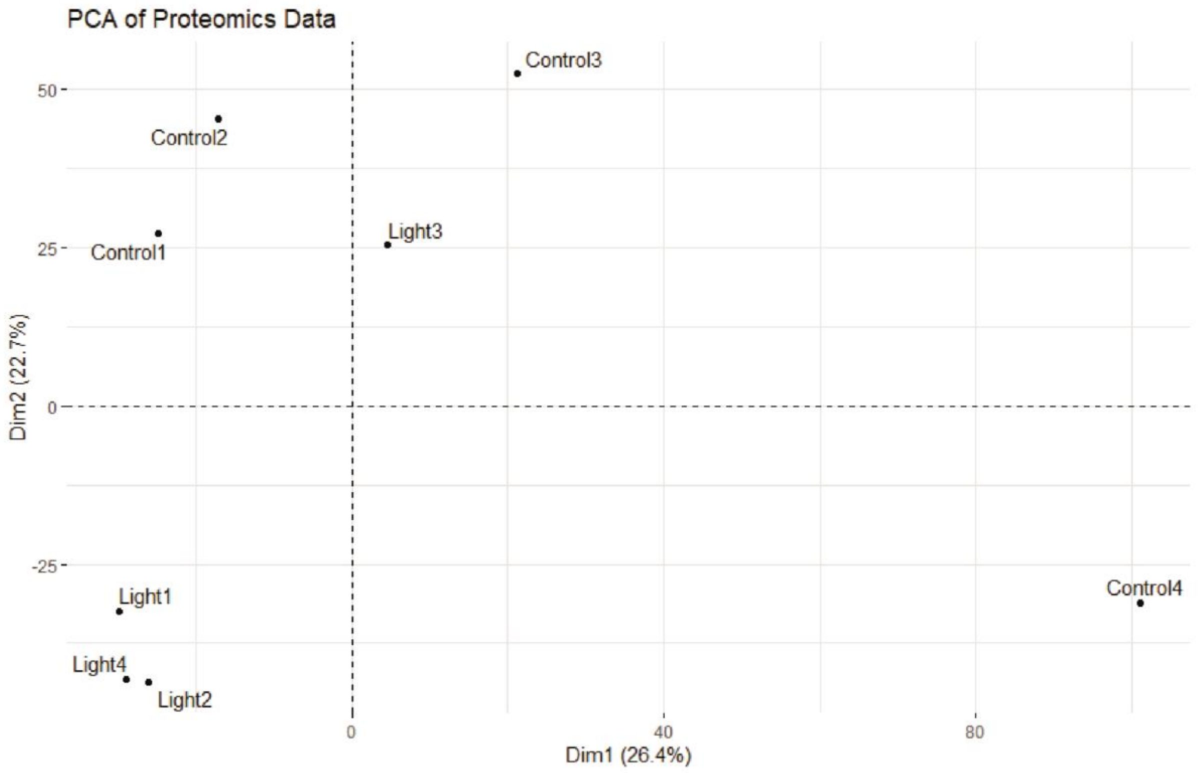


**Supplementary Figure 2. Principal component analysis (PCA) of DIV210 control and light-conditioned ROs submitted for proteomics.**

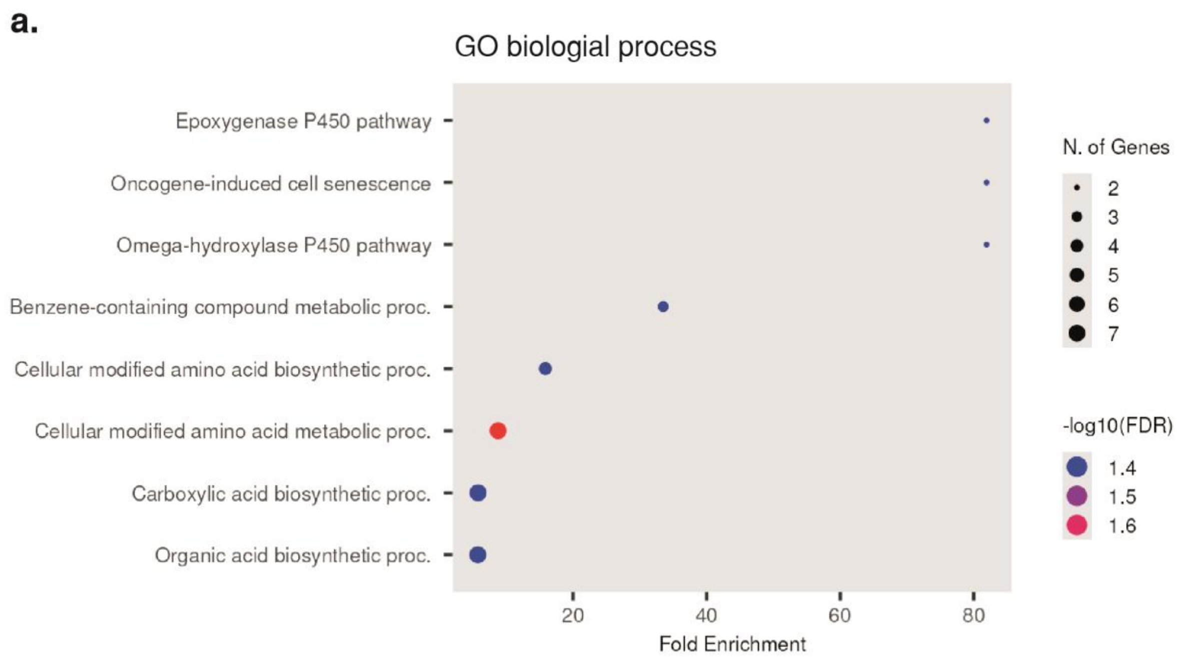


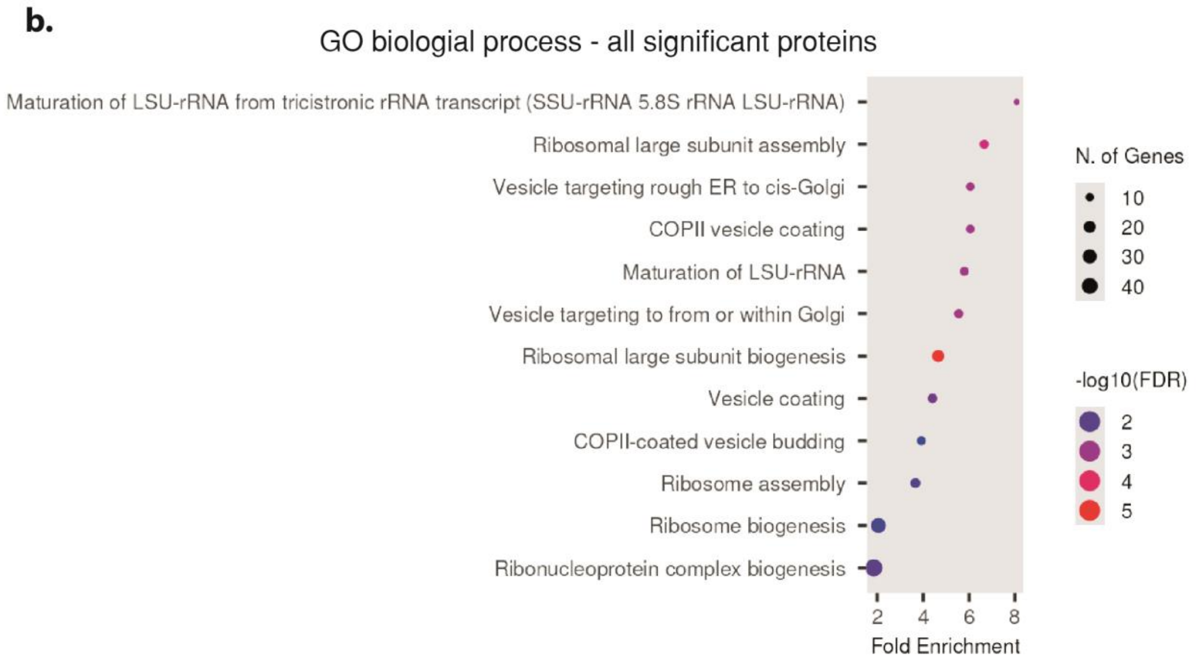


**Supplementary Figure 3. Gene ontology (GO) analysis of proteomics screen. (a)** GO biological process for proteins with p-value < 0.05 and Log2FC > ± 0.5. **(b)** GO biological process for proteins with p-value < 0.05, regardless of the Log2FC. For both analyses, all detected proteins were used as background, and a false discovery rate cut-off value of 0.05 was used.

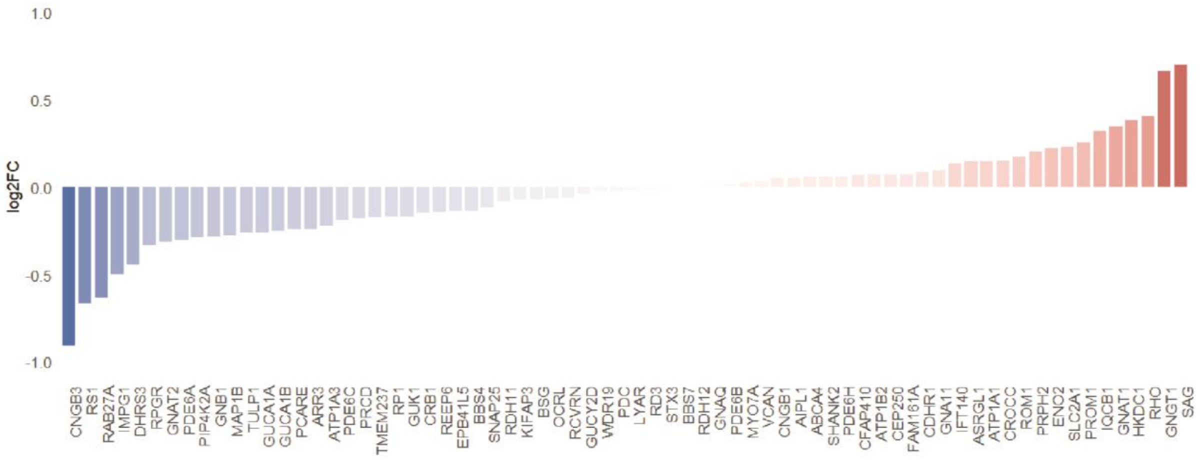


**Supplementary Figure 4. Photoreceptor inner segment and photoreceptor outer segment protein expression.** Protein markers were based on GO annotations for photoreceptor inner segments (GO:0001917) and photoreceptor outer segments (GO:0001750). Data represents the log2FC of light-conditioned ROs compared to control ROs.
